# Mitochondria alterations in fibroblasts from sporadic Alzheimer’s disease (AD) patients correlate with AD-related clinical hallmarks

**DOI:** 10.1101/2023.12.21.570579

**Authors:** Fanny Eysert, Paula-Fernanda Kinoshita, Julien Lagarde, Sandra Lacas-Gervais, Laura Xicota, Guillaume Dorothée, Michel Bottlaender, Frédéric Checler, Marie-Claude Potier, Marie Sarazin, Mounia Chami

## Abstract

**INTRODUCTION:** Mitochondria dysfunctions are key features in Alzheimer’s disease (AD). The occurrence of these disturbances in AD patient’s peripheral cells and their potential correlation with disease progression are under investigated.

**METHODS:** We studied mitochondrial structure, function and mitophagy in fibroblasts including healthy volunteers and AD patients at the prodromal (AD-MCI) or demented (AD-D) stages. We carried out correlation studies with clinical cognitive scores, Aβ plaques burden and the accumulation of peripheral amyloid precursor protein C-terminal fragments (APP-CTFs).

**RESULTS:** We unveiled progressive alteration of mitochondria structure as well as specific mitochondrial dysfunctions signatures in AD-MCI and AD-D fibroblasts and show defective mitophagy and autophagy linked to impaired lysosomal activity in AD-D fibroblasts. We reported on significant correlations of a subset of these dysfunctions with cognitive decline, AD-related clinical hallmarks and peripheral APP-CTFs accumulation.

**DISCUSSION:** This study emphasizes the potential use of peripheral cells for investigating AD pathophysiology and likely diagnosis.

## 1- BACKGROUND

Mitochondria studies in the field of Alzheimer’s disease (AD) identified novel pathophysiological mechanisms accounting for the disease development. It is now recognized that mitochondrial dysfunctions contribute to neurodegeneration occurring in AD^1,2^. Indeed, mitochondrial dysfunctions have been observed in post-mortem human brain samples and were demonstrated in AD cellular models as well as in the brain of AD mice^1^. These alterations include impaired mitochondrial membrane potential (ΔΨm), reduced ATP production, increased mitochondrial reactive oxygen species (mitROS) levels and altered mitochondrial fission and fusion balance^3,4^. Moreover, the specific process that removes damaged mitochondria called mitophagy is altered in AD^5,6^. The accumulation of the damaged mitochondria is a source of noxious material (e.g. toxic mitROS) likely contributing to a deleterious vicious cycle driving the disease progression.

In AD transgenic mice brain, mitochondrial dysfunctions and mitophagy defect occurred early since they are observed before the appearance of the extracellular deposits of the β-amyloid peptides (Aβ) in senile plaques^3^. These Aβ peptides are among the products of the cleavage of the transmembrane amyloid precursor protein (APP). In the amyloidogenic pathway, APP is first cleaved by the β-secretase generating APP C-terminal fragment (APP-CTF) of 99 amino acids (C99), that is further cleaved by the γ-secretase to produce Aβ peptides and the amyloid precursor protein intracellular domain (AICD). Alternatively, APP and C99 can also be cleaved by the α-secretase producing the APP-CTF of 83 amino acids (C83)^7^. We recently demonstrated that the accumulation of APP-CTFs (C99 and C83) in cellular models triggers, in an Aβ-independent manner, excessive mitochondrial morphology alteration as well as enhanced mitROS production and mitophagy impairment^5^. We further demonstrated alterations of mitochondrial structure and mitophagy process in the brain of transgenic mice accumulating APP-CTFs and strengthened these observations in human post-mortem sporadic AD brains^5^. Accordingly, defects of the general autophagy and lysosomal activity in AD have been associated with APP-CTFs accumulation^8–10^.

Mitochondrial dysfunctions in AD are not restricted to the brain. Indeed, several studies reported an alteration of mitochondria structure and function in peripheral cells from AD patients such as lymphocytes, peripheral blood mononuclear cells (PBMCs) or fibroblasts^11–13^. Other studies have unraveled that mitochondrial dysfunctions occurred in peripheral cells of familial forms of the disease (FAD)^14,15^ representing 1% of AD cases and linked to mutations in APP or presenilin 1 and 2 genes (coding for the catalytic core of γ-secretase), as well as in sporadic AD (SAD)^12^ with a non-Mendelian (complex) transmission and representing the majority of AD cases^16^. It remains however uncertain if these mitochondrial dysfunctions taking place in AD peripheral cells occurred at the prodromal stage and could thus be considered as potential early hallmarks correlating with clinical severity or pathophysiological AD markers.

We studied herein fibroblasts obtained from a well characterized cohort of healthy volunteers and SAD patients at mild cognitive impairment (AD-MCI) or dementia (AD-D) stages ^17,18^ and investigated several aspects of mitochondrial structure and function as well as the key steps underlying mitophagy and autophagy process. The originality of the current study also stands in the correlative analyses between the observed alterations and the clinical data.

## 2. METHODS

### 2.1- Patients and control individuals

Human primary fibroblasts were obtained from the IMABio3 and Shatau7-Imatau study cohorts (NCT01775696 and NCT02576821-EudraCT2015-000257-20) (CTRL n=9; AD-MCI n=11; AD-D n=9), approved by a French Ethics Committee. All subjects provided written informed consent prior to participating.

Patients with AD were included according to the following criteria: (i) cognitive impairment characterized by a predominant progressive episodic memory deficit; (ii) positive pathophysiological markers of AD, defined by CSF AD profile and amyloid Pittsburgh compound B (PiB)-PET imaging. Controls were recruited according to the following criteria: (i) Mini-Mental State Examination (MMSE) score ≥ 27/30 and normal neuropsychological assessment; (ii) Clinical Dementia Rating-Scale Sum of Boxes (CDR-SOB) equal to 0; (iii) no history of neurological or psychiatric disorders; (iv) no memory complaint or cognitive deficit, and (v) amyloid PiB-PET imaging < 1.5 except for one individual who a PiB-GCI value of 2.18 but a CDR-SOB equal to 0 and a MMSE at 30 at inclusion date and was clinically stable over two years after inclusion. We did not include subjects with severe cortical or subcortical vascular lesions, a history of autoimmune and inflammatory diseases or psychiatric disorders and suspicion of alcohol or drugs abuse. Participants followed a complete clinical and neuropsychological assessment, and ^11^C-PiB PET imaging as previously described^18,19^, and were followed up annually for 2 years. Skin biopsies were obtained during the 1-year or final (2-year) visit. Demographic description of the sample according to diagnostic group are described in Table 1.

**Table 1:**
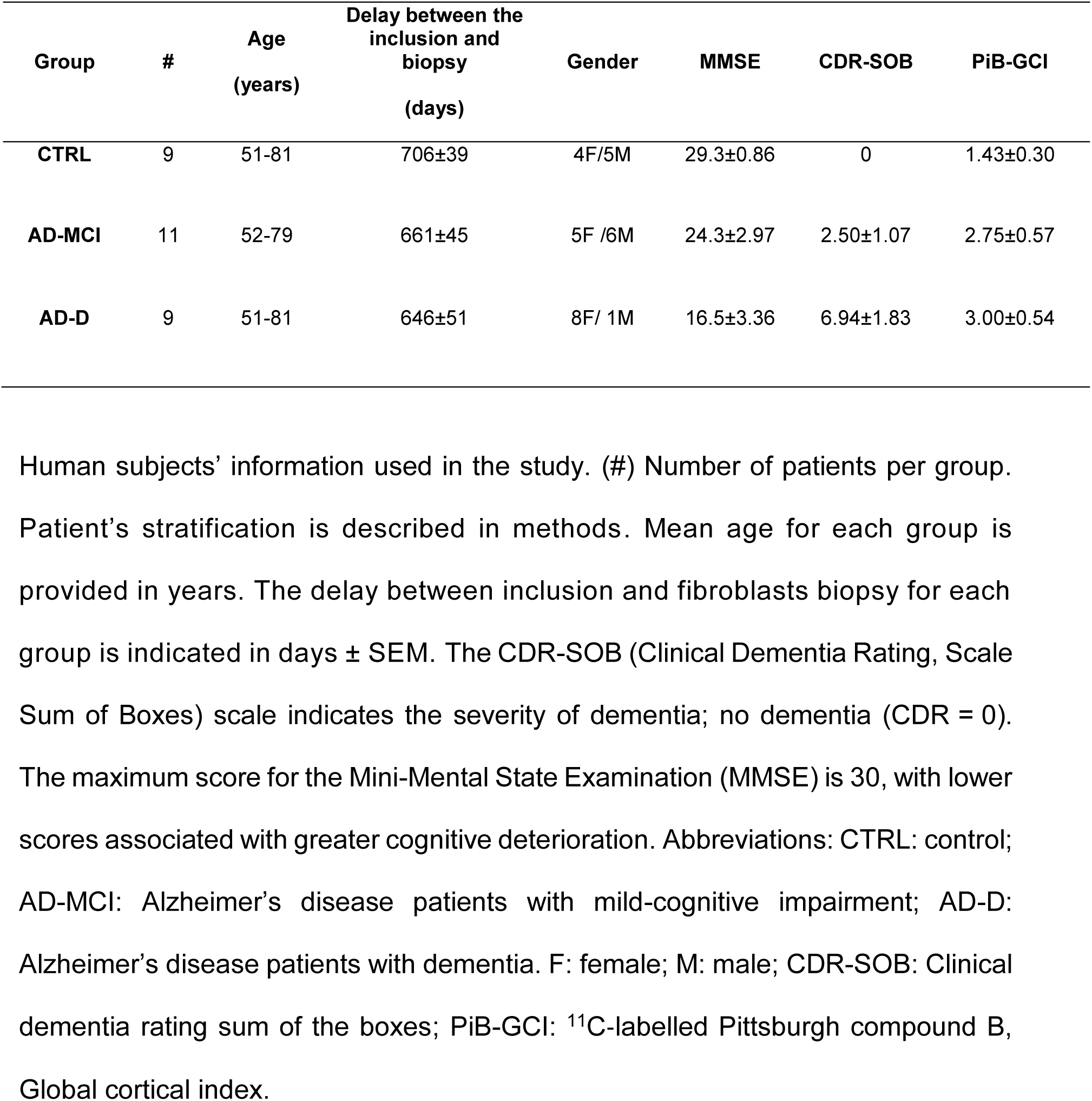
Demographic characteristics of the fibroblasts cohorts at patients inclusion.

### 2.2- Primary fibroblasts culture

Fibroblasts were cultured as described previously^20^ and kept under liquid nitrogen at the DNA & Cell Bank core facilities of the Brain and Spinal Cord Institute (ICM). Briefly, cells were cultured at 37°C in a humidified atmosphere containing 5% CO2 in DMEM (Dulbecco modified Eagle’s minimal essential medium) supplemented with 10% heat-inactivated fetal bovine serum (FBS), 1% pyruvate sodium, Penicillin (100 U/mL) and streptomycin (50 μg/mL). All experiments were performed using cells between the 4^th^ and the 12^th^ passages. Experiments were performed on batches from the same passages.

### 2.3- Chemicals and antibodies

When indicated, fibroblasts were treated with the respiratory chain uncoupling agents, Carbonyl cyanide 3-chlorophenylhydrazone (CCCP) at 10µM (Millipore-Sigma, C2759) for 6 hours, or with Trifluoromethoxy carbonylcyanide phenylhydrazone, Carbonyl cyanide 4-(trifluoromethoxy) phenylhydrazone (FCCP) at 10 µM (Millipore-Sigma, C2920) for 1 min. Fibroblasts were also treated with oligomycin at 10µM (Abcam, 141829) and antimycin A at 1µM (Sigma, A8674) overnight.

We used the following commercially available antibodies: β-actin (A5316, Sigma-Aldrich), APP-Cter (A8717, Gift from P. Fraser, Toronto), Chaperonin 10 (CPN10) (ADI-SPA-110, Enzo Life Sciences), CoxII (12C4F12, ThermoFisher Scientific), and TOMM20 (612278, BD Transduction Laboratories). The secondary antibodies used for western blot studies were goat anti-mouse and goat anti-rabbit (115-036-003 and 111-036-045 respectively, Jackson Immuno Research).

### 2.4- Mitochondrial fraction preparation and SDS page analysis

Cells growing were harvested with 0,05% trypsin (Gibco), washed with PBS, sedimented by centrifugation and resuspended in isolation buffer (250 mM D-Mannitol, 5 mM HEPES pH 7.4, 0.5 mM EGTA, and 0.1% BSA) supplemented with protease inhibitor mixture (Sigma, P2714-1BTL). After 15 min of incubation on ice, cells were disrupted by 100 strokes of a glass Dounce homogenizer. The lysates were centrifuged at 2500 × *g* at 4 °C for 5 min to remove unbroken cells and nuclei. The supernatant was centrifuged at 10,000 × *g* at 4 °C for 10 min to pellet mitochondrial fraction which was suspended in isolation buffer supplemented with protease inhibitors. The protein concentration was determined according to the Bradford method (BioRad protein Assay, Spectrophotemeter Eppendorf biophotometer plus at 595 nm). Full-length APP and APP-CTFs were resolved on 16.5% Tris-Tricine SDS-PAGE then transferred onto nitrocellulose membranes which were boiled in PBS. All the other proteins were resolved on 16% Tris-Glycine SDS-PAGE. Then, membranes were saturated in 5% skimmed milk TBS tween buffer (TBS-T), and incubated overnight at 4°C with specific primary antibodies. After washing with TBS-T, membranes were incubated with HRP-conjugated antibodies (Jackson ImmunoResearch, 1/5000) for 1 hour at room temperature. Membranes were rinsed with TBS-T and visualized using ImageQuant LAS4000.

### 2.5- Mitophagy and autophagy analyses

Twenty-four hours after seeding on glass coverslips, cells were transiently transfected with LC3-GFP (Addgene plasmid #11546), LAMP1-GFP probe (Addgene plasmid #34831)^21^, the mitochondrial Mit-RFP probe^22^ and the Cox8-EGFP-mCherry mitophagy reporter (a gift from David Chan; Addgene plasmid # 78520)^23^ using jetPRIME (polyplus transfection, #114-15) according to the manufacturer’s instructions.

Cells transfected with Mit-RFP probe were treated 24 hours later with drugs when indicated and fixed with 4% paraformaldehyde for 15 min at room temperature. Other cells were incubated 30 min with Lysotracker-Red DND-99 (1:20,000, Invitrogen) at 37°C and fixed with PFA 4 %. Cells were rinsed with PBS and stained with DAPI (1:10,000, Roche) before coverslips mounted on glass slides with Vectamount medium (Vector).

The Cox8-EGFP-mCherry mitophagy reporter is based on differences in pKa of green fluorescent protein (EGFP), and mCherry expressed in tandem with the mitochondrial localization signal of Cox8. In mitophagy, mitochondria are delivered to lysosomes where the low pH (pH=4) quenches the EGFP signal. The result is that a portion of mitochondria fluoresce red only. For Cox8-EGFP-mCherry experiments, images were acquired by live imaging using Zeiss LSM 780 with 63X Objective. Data are presented as the percentage of cells undergoing mitophagy or as the number of red dots per cell. A threshold of a single or more red-alone puncta per cell was applied to all cells expressing Cox8-EGFP-mCherry probe. Fixed cells were visualized using Leica SP5 microscope with 63X objective. The colocalization quantification was determined using Fiji plug-in JACoP (Just Another Colocalization Plug-in^24^). The quantifications of LC3 and LAMP1 dots size and count and LysoTracker intensity were assessed using Analyze particles function of Fiji software. Mitochondrial network volume was quantified thanks to 3D object counter plugin of Image J and mitochondria branches length and number as well as end-points and junction number were quantified using skeletonize plugin of Image J^25^. The quantification was obtained from at least 25 cells per patient.

### 2.6- In vitro Cathepsin D activity assay

CTRL, AD-MCI and AD-D fibroblasts were harvested with 5 mM PBS-EDTA, centrifuged at 600 x *g* for 5 min, washed in PBS and centrifuged again at 600 x *g* for 5 min. The pellet was resuspended in Tris–HCl (10 mM, pH 7.5). To monitor cathepsin D (CTSD) activity, 25 μg of protein extracts were incubated in acetate buffer (25 mM, pH 4.5, and 8 mM L-cysteine HCl) containing CTSD substrate (Enzo BML-P145-001, 50µM) in the absence or presence of pepstatin A (20 μM, Sigma). CTSD activity corresponds to the pepstatin A-sensitive fluorescence recorded at 320 nm (excitation) and 420 nm (emission) using a fluorescence plate reader (Varioskan, ThermoFisher scientific). The fluorescence was recorded every 30 seconds during 45 min and the CTSD activity was calculated as the area under the curve (AUC) and the slope in the linear range corresponding to the initial 5 min.

### 2.7- Mitochondrial potential and mitochondrial superoxide measurements

Tetramethyl rhodamine methyl ester (TMRM) (ThermoFisher, #T668) is a cell-permanent dye that accumulates in active mitochondria with intact membrane potentials. Loss of mitochondrial membrane potential indicating mitochondrial membrane depolarization triggers reduced TMRM accumulation. On the contrary, elevated TMRM intensity reflects an increase in mitochondrial membrane potential, indicating mitochondrial membrane hyperpolarization^26^. MitoSOX Red (Invitogen, M36008) is a fluorogenic dye for highly selective detection of superoxide in the mitochondria of living cells^27^. Cells were rinsed with PBS, and incubated in TMRM at 2 nM or MitoSOX at 5 µM in DMEM for 30 min, at 37 °C. Cells were imaged directly (TMRM staining) or rinsed with KRB buffer (135mM NaCl, 5mM KCl, 1mM MgSO4, 0.4 mM K2HPO4, 20 mM HEPES, 0.05 mM CaCl2, 1g/L Glucose, pH 7.4) (MitoSOX staining). Images were acquired by live imaging using Zeiss LSM 780 with 63X Objective. Both TMRM and MitoSOX fluorescence median intensities were analyzed using Fiji software. The quantification was obtained from at least 25 cells per patient.

### 2.8- Seahorse Mito stress test and ATP rate analyses

Measurements of aerobic respiration and glycolysis were conducted with the Seahorse Bioscience XFe96 bioanalyzer using the seahorse XF Mito Stress Test Kit (Agilent #103015-100) and the Seahorse XF Real-Time ATP Rate Assay Kit (Agilent #103592-100). Fifteen thousand cells per well were seeded on XFe96 cell culture microplates (Agilent #102416-100) one day before the experiment. On the day of the experiment, the culture medium was replaced with Seahorse Base Medium (Agilent #103334-100) supplemented with 1mM pyruvate, 2 mM glutamine and 10 mM glucose, and incubated for 30 min at 37°C in a CO_2_-free incubator. For the Seahorse XF Real-Time ATP Rate Assay, the medium was replaced a second time after the incubation time, before loading the cell culture microplate into the Seahorse Analyzer. After measuring basal respiration, oligomycin (5µM), FCCP (2µM), and rotenone/antimycin A (0,5 µM each) were added in a sequential order to each well for XF Mito Stress Test, while oligomycin and rotenone/antimycin A were added for Real-Time ATP Rate Assay. Data were analyzed using the XF Cell Mito Stress Test or XF Cell Real-Time ATP Rate Report Generator respectively. After the assay, fibroblasts were fixed with PFA 4 % and stained with DAPI (Roche; 1:10,000) for 5 min. Cytation 5 Biotek was then used to count the number of cell nuclei (cell number) in each well. The normalization of the experiments is based on the relative cell number per well.

### 2.9- Transmission electron microscopy

For ultrastructure analysis, cells were fixed in 1.6% glutaraldehyde in 0.1 M phosphate buffer (pH 7.4), rinsed with cacodylate buffer 0.1 M, and then post-fixed in osmium tetroxide (1% in cacodylate buffer) reduced with potassium ferrycyanide (1%) for 1 hour. Cells were dehydrated with several incubations in increasing concentrations of ethanol or acetone, respectively, and embedded in epoxy resin (EPON), and 70 nm ultrathin sections were contrasted with uranyl acetate and lead citrate and observed with a Transmission Electron Microscope (JEOL JEM 1400) operating at 100 kV and equipped with Olympus SIS MORADA camera. We used Fiji software to analyze mitochondria ultrastructure. Images were taken in blind. The quantification of mitochondrial ultrastructure by transmission electron microscopy was done in at least 15 different fields (150-350 mitochondria per patient). The quantification of mitochondria classes was first done from individual images from the same patient and the means +/- SEM of each mitochondria class was then reported for each fibroblast group.

### 2.10- Statistical analyses

Data were expressed as means ± SEM. Sample size for each experiment is indicated in the figure captions. Data were analyzed with GraphPad Prism version 8 for Windows (GraphPad Software, La Jolla, CA, USA). We used Kruskal–Wallis test and Dunn’s multiple comparisons post-test unless otherwise indicated. Correlation analyses were performed by linear regression used to determine *P* and goodness of fit (R^2^) values taking into consideration the age of patients as well as the delay between the date of inclusion and the date of skin biopsy as covariates using InVivoStat software. Significant differences are: \**P* < .05, \*\**P* < .01, \*\*\**P* < .001, \*\*\*\**P* < .0001 and ns: not significant.

## 3 RESULTS

### 3.1- Mitochondrial ultrastructure is altered in AD-D fibroblasts and is associated with cognitive impairments

We assessed mitochondria ultrastructure by transmission electron microscopy and revealed that mitochondria number is not different between fibroblasts isolated from CTRL individuals, AD-MCI and AD-D (Figure 1A, D). We classified mitochondria ultrastructure in four classes based on the organization of the cristae and the density of the matrix as previously described^5^ (class I: healthy dark mitochondria with uniform matrix filled with dense regular distributed cristae; class II: mitochondria with disrupted cristae and loss of matrix density; class III: empty mitochondria with disorganized cristae; and class IV: swollen mitochondria with disorganized cristae) (Figure 1A, B). This classification unveiled a majority of healthy class I mitochondria (84.13%) and a minor proportion of class II (8.51%), class III (2.70%) and class IV (4.65%) mitochondria in CTRL fibroblasts (Figure 1C, D). However, we reported that the distribution of mitochondria sub-types is modified in AD fibroblasts. Thus, AD-MCI and AD-D fibroblasts displayed lower healthy class I mitochondria (75.27% and 61.48% respectively) and an enhancement of class II (9.59% and 17.12%, respectively), class III (8.21% and 10.61%, respectively) and class IV (6.92% and 10.78% respectively) mitochondria as compared to CTRL cells (Figure 1C, D). Importantly, the drop in the percentage of mitochondria class I in AD-D fibroblasts is statistically significant as compared to that in CTRL fibroblasts.

**Fig. 1.**
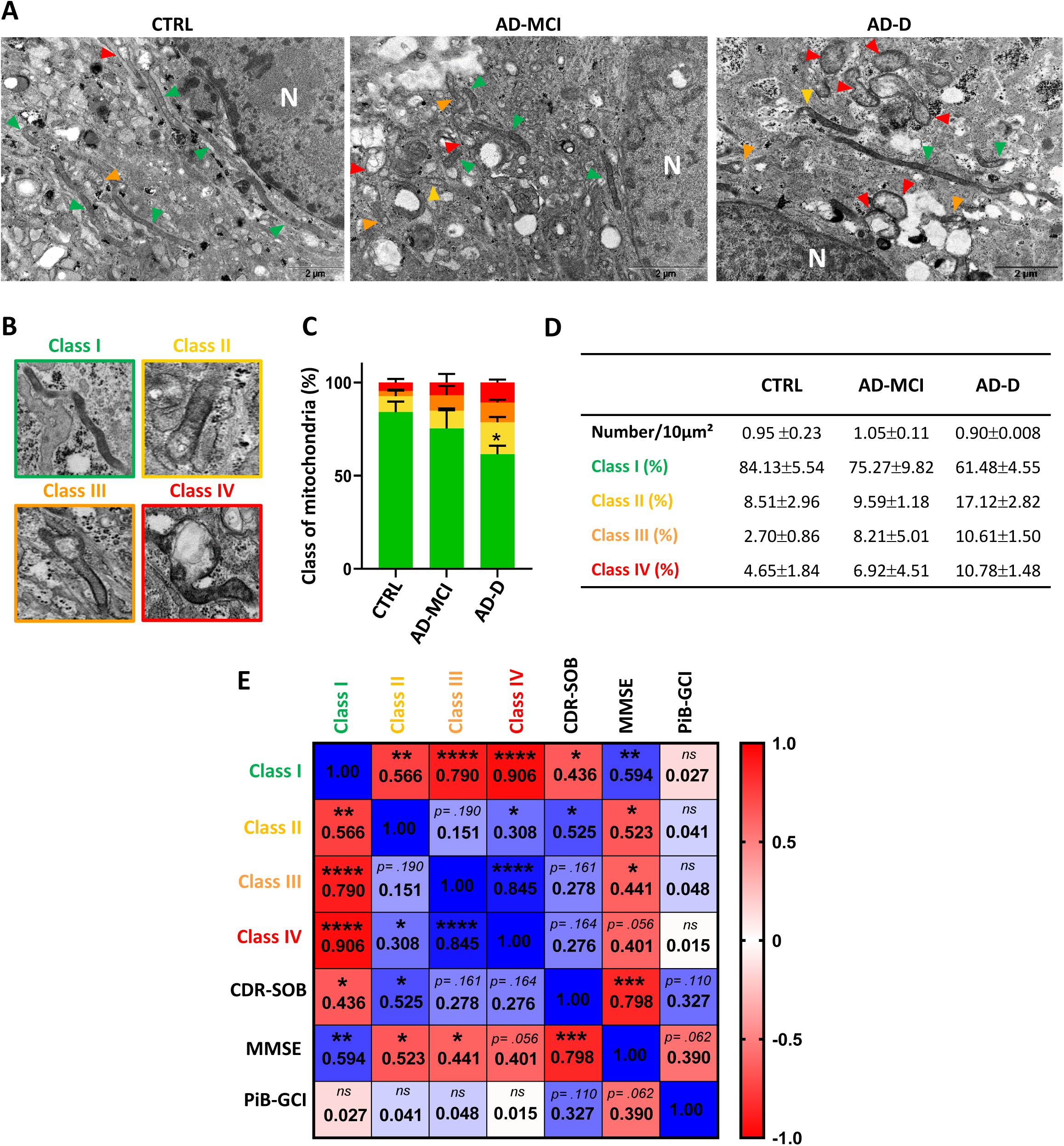
Mitochondrial classes distribution is different between CTRL, AD-MCI and AD-D fibroblasts and correlates with cognitive impairments. **A)** Representative electron microscopy ultrastructure in CTRL, AD-MCI and AD-D fibroblasts. N: nuclei. Colored arrowheads indicate mitochondria (Green = Class I, Yellow= Class II, Orange= Class III and Red=Class IV). Scale bar=2µm. **B)** Representative images of mitochondria classes I, II, III, and IV. **C)** Quantitative graph of mitochondria classes distribution ± SEM. * p=.013 using 2 ways ANOVA and Tukeys multiple comparisons test versus CTRL class I mitochondria. **D)** Table presenting the average number ± SEM of mitochondria per 10µm² and the percentage of mitochondria classes ± SEM obtained from CTRL (n=4), AD-MCI (n=4) and AD-D (n=5) fibroblasts. The quantification was done in at least 15 different fields (between 150-350 mitochondria/patient). **E)** Heat map of the correlation matrix computing linear regression value (R²) between mitochondria classes (I, II, III and IV), and the clinical Dementia Rating (Sum Of Boxes) (CDR-SOB), the Mini Mental State Examination (MMSE) and the β-amyloid plaques burden assessed using the ^11^C-labelled Pittsburgh compound B (Global Cortical Index) (PiB-GCI) through positrons emission tomography (PET-scan) at patient’s inclusion. Color scale of 1 corresponds to the maximum positive correlation value (blue) and −1 corresponds to the maximum negative correlation value (red). Correlation analyses took into consideration the age of patients and the delay between the inclusion date and the date of skin biopsy as covariates. *p<.05; **p<.01; ***p<.001; ****p<.0001 for the correlation of every pair of data sets.

The correlation analyses including CTRL, AD-MCI and AD-D fibroblasts and integrating mitochondria classes, CDR-SOB and MMSE scores and PiB-GCI at inclusion and taking into consideration the age of patients as well as the delay between the date of inclusion and the date of skin biopsy as covariates revealed as expected: i) significant negative correlations between healthy (class I) and damaged mitochondria (classes II, III, and IV) (Figure 1E); and, ii) a significant inverse correlation between the CDR-SOB score and the MMSE score (Figure 1E, and Table 1). Importantly, we demonstrated that MMSE score has a significant positive correlation with healthy mitochondria (class I) and significant negative correlation with damaged mitochondria (class II, III and IV) (Figure 1E). In parallel, we reported that the CDR-SOB score has significant negative correlation with healthy mitochondria (class I), and positively correlate with dysfunctional mitochondria reaching statistical significance with mitochondria class II (Figure 1E). However, these mitochondria ultrastructure alterations did not correlate with PiB-GCI (Figure 1E), Tau, p-Tau, or Aβ42 levels in CSF (data not shown).

Altogether, these data demonstrate a relevant alteration of mitochondria ultrastructure in AD-D versus CTRL fibroblasts. They also point out that the alteration of mitochondrial ultrastructure is closely associated with cognitive impairment (CDR-SOB and MMSE scores).

### 3.2- Mitochondrial volume and network organization is impacted in AD-D fibroblasts

We then analyzed in depth mitochondrial morphology using confocal imaging and three-dimensional (3D) reconstitution of mitochondria network in fibroblasts transiently transfected with Mit-RFP probe (Figure 2A). In order to investigate the potential differences between AD-MCI, AD-D and CTRL fibroblasts towards mitochondrial stress conditions, these analyses were performed under basal condition and upon CCCP treatment known to trigger mitochondrial membrane depolarization and mitochondrial network fragmentation. As expected, CCCP treatment decreases the mitochondrial volume in the three groups (Figure 2B). Interestingly, we reported a tendency but not significant decrease of the mitochondrial volume in AD-D fibroblasts as compared to CTRL and AD-MCI cells under basal condition. These results suggest a trend towards fragmentation of the mitochondrial network in AD-D fibroblasts as compared to AD-MCI and CTRL cells.

**Fig. 2.**
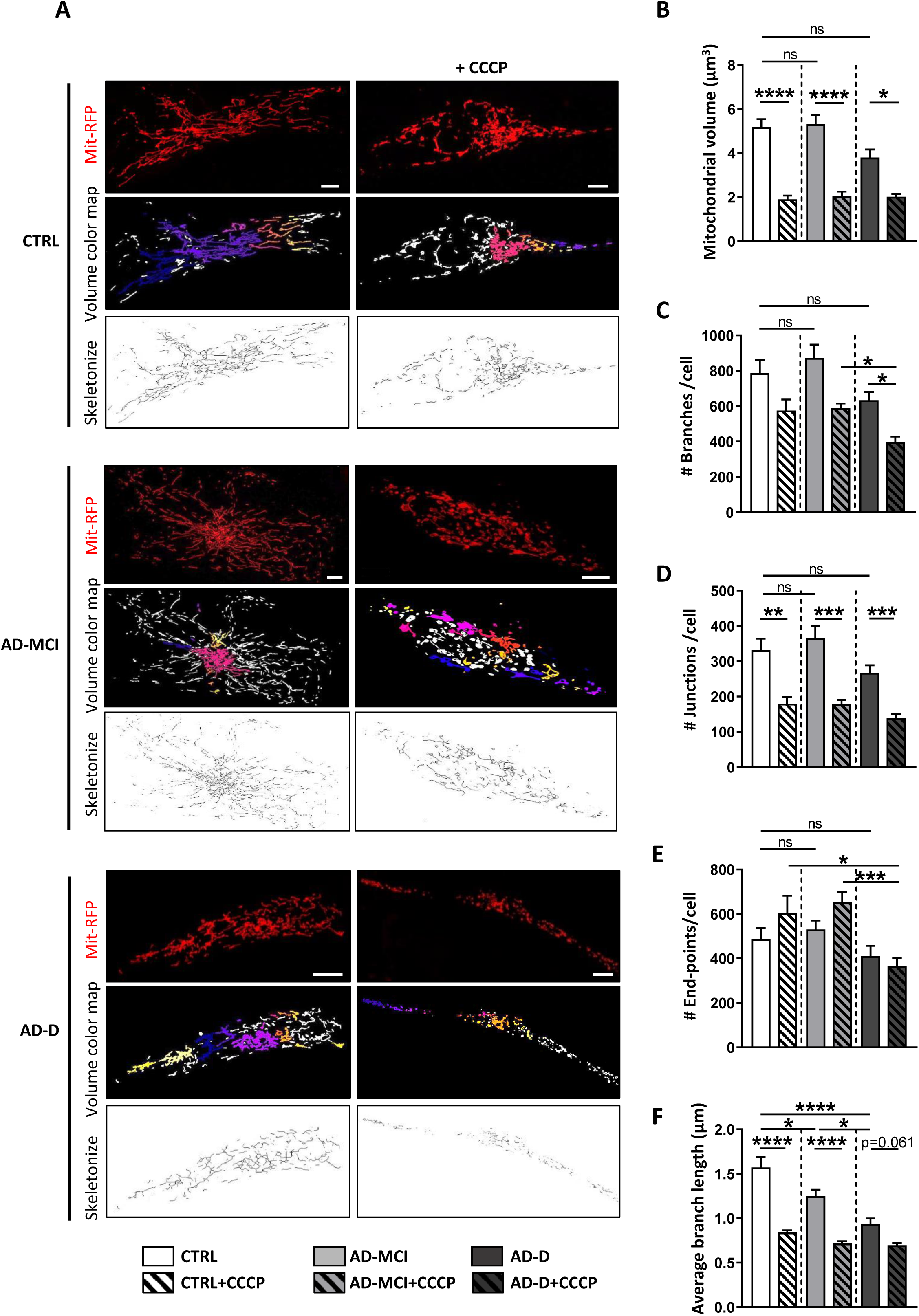
Mitochondrial volume and network organization is more impacted in AD-D versus AD-MCI fibroblasts. **A)** Representative images of mitochondrial network morphology in CTRL, AD-MCI and AD-D fibroblasts transfected with Mit-RFP probe and treated or not with 10µM CCCP for 6 hours. Scale bar=2µm. Original images were processed with Fiji 3D Counter plugin providing the volume color map and skeletonize morphology. **B-F)** Quantitative graphs of the mitochondrial volume (µm^3^) (B), the average network branches per single cell (C), the mean branch-length (µm) in mitochondrial network in individual cells (D), the number of junctions (E), and the number of end-points per single cell (F). Means ± SEM were obtained from CTRL (n=4), AD-MCI (n=4) and AD-D (n=4). *p<0.05; **p< .01; ***p< .001; ****p< .0001.

To fully demonstrate this assumption, we evaluated several mitochondrial structure features including the average number of branches (Figure 2C), junctions (Figure 2D) and end-points (Figure 2E), as well as the average of branch-length in mitochondrial network per single cell (Figure 2F). Consistent with mitochondrial volume analyses (Figure 2B), our data indicated no significant modifications in the number of branches, junctions or end-points between AD-MCI versus CTRL fibroblasts under basal condition (Figure 2C-E), and showed a trend diminution of these parameters in AD-D fibroblasts (Figure 2C-E). Specifically, CCCP treatment significantly reduced the number of junctions in the three groups of cells (Figure 2D), and revealed greater sensitivity of AD-D fibroblasts versus CTRL and AD-MCI branches number (Figure 2C) and end-points number (Figure 2E). Importantly, we noticed a significant diminution of the mitochondria average branch-length network in AD-MCI and AD-D cells as compared to CTRL (Figure 2F) and also between AD-MCI and AD-D cells (Figure 2F). In addition, CCCP treatment drastically reduced the average branch-length in CTRL and AD-MCI to reach that observed in AD-D fibroblasts (Figure 2F). Together, these analyses demonstrate a progressive mitochondrial network alteration in AD-D fibroblasts versus AD-MCI and CTRL fibroblasts.

### 3.3- Mitochondria function and -related metabolic activities are dysfunctional in AD-MCI and AD-D fibroblasts

As mitochondria structure is altered in AD-MCI and AD-D fibroblasts, we hypothesized that mitochondrial function could also be impaired in these cells versus CTRL ones. Mitochondrial function is classically defined by the membrane potential as well as ROS production. First, we assessed mitROS level by using MitoSox probe and confocal microscopy analyses. As for mitochondrial structure analyses, these experiments were performed under basal condition and mitochondria dysfunction triggered by oligomycin and antimycin A (OA) known to block respectively mitochondrial ATPase and the respiratory chain complex III. We first showed a significant increase in Mitosox intensity in AD-MCI as compared to CTRL fibroblasts (Figure 3A, B). Unexpectedly, we noticed a non-significant decrease in mitROS level in AD-D fibroblasts as compared to CTRL that reached statistical significance versus AD-MCI fibroblasts (Figure 3B). However, when fibroblasts were challenged with OA, MitoSox intensity was drastically increased in AD-D fibroblasts as compared to non-treated AD-D fibroblasts and to CTRL fibroblasts treated with OA, indicating enhanced sensitivity of AD-D fibroblasts towards OA treatment (Figure 3B).

**Fig. 3.**
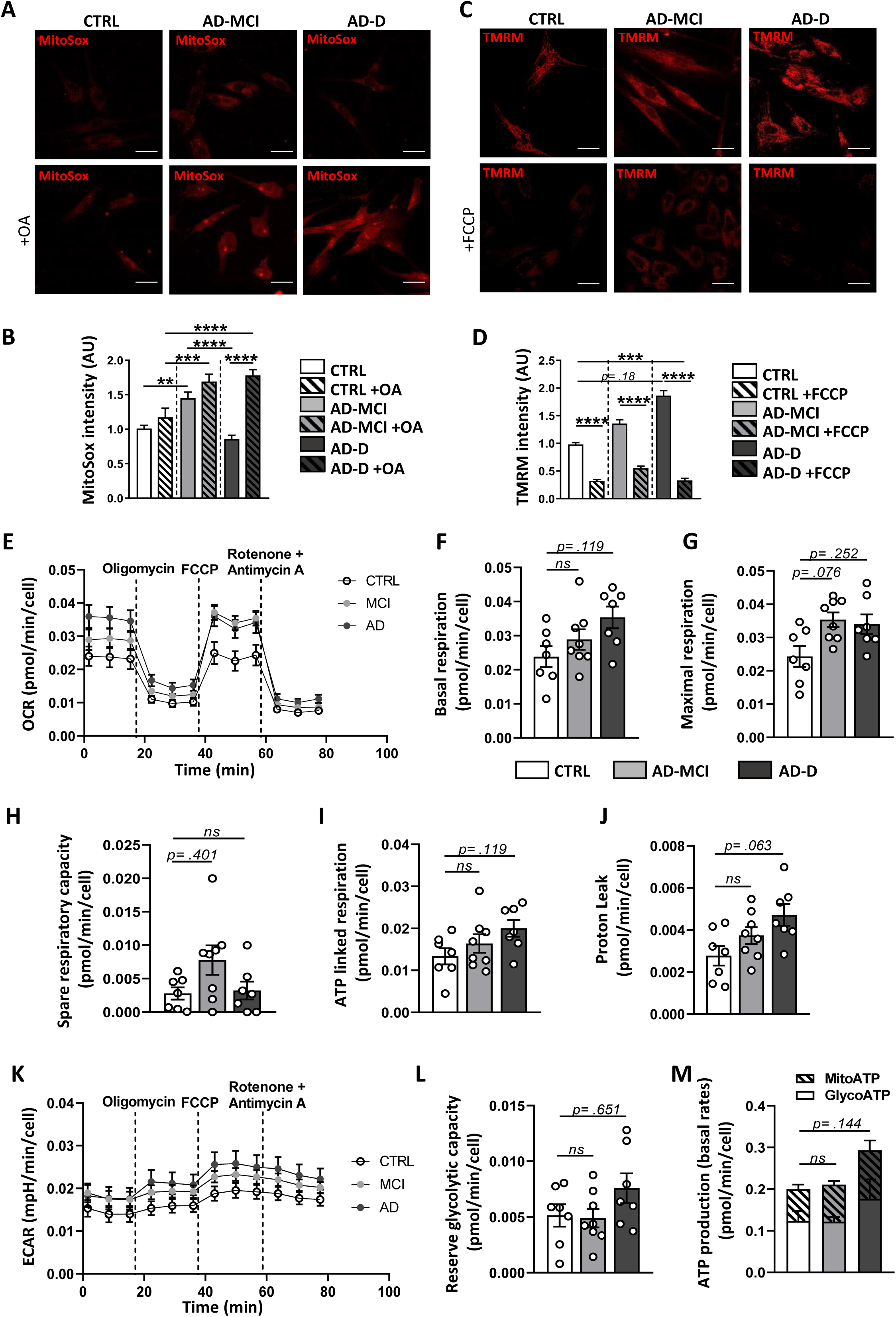
Mitochondrial ROS, membrane potential and metabolism are altered in a disease-specific manner in AD-MCI in AD-D fibroblasts. **A)** Representative confocal microscopy images obtained with MitoSox probe. **B)** Quantitative graph of Mitosox mean intensity ± SEM obtained from CTRL (n=4), AD-MCI (n=4) and AD-D (n=4). **C)** Representative confocal microscopy images obtained with TMRM probe. **D)** Quantitative graph of TMRM mean intensity ± SEM obtained from CTRL (n=3), AD-MCI (n=3) and AD-D (n=3). **A, C)** Scale bar= 10µm. **B, D)** **p<0.01; ***p<0,001; ****p<0.0001. **E-L)** Cell respiratory capacity using SeaHorse XFe96 extracellular flux analyser (Mito Stress Test) obtained in CTRL (n= 7), AD-MCI (n=8) and AD-D (n=7) fibroblasts. **M)** ATP production measured using SeaHorse XFe96 extracellular flux analyser (ATP Rate Assay) obtained in CTRL (n= 5), AD-MCI (n=6) and AD-D (n=5) fibroblasts. Data are expressed in pmol/min/cell ± SEM or mpH/min/cell ± SEM obtained from 5 replicates for each type of fibroblasts. ns: not significant using 2way ANOVA test.

We analyzed mitochondrial membrane potential under basal condition and upon treatment with FCCP using TMRM probe and confocal microscopy (Figure 3A, D). As expected, fibroblasts challenged with FCCP treatment showed a drastic mitochondrial membrane depolarization in the three groups of cells (Figure 3D). Moreover, we observed a gradual upward of TMRM intensity in AD-MCI and AD-D fibroblasts as compared to CTRL (Figure 3C, D). However, we did not reveal any significant correlations between TMRM or MitoSox intensities and clinical data (CDR-SOB, MMSE) or the β-amyloid plaques burden (PiB-GCI) (Suppl. Fig. 1A-F).

To characterize the metabolic constants in the IMABio3 fibroblasts, we measured the oxygen consumption rate (OCR) and the extracellular acidification rate (ECAR) using Seahorse technology and a set of mitochondrial respiration modulators (Oligomycin, FCCP, and Rotenone/Antimycin A) (Figure 3E, K). This protocol allowed us to determine basal respiration, maximal respiration, spare respiratory capacity, oxygen consumption associated with ATP synthesis, and oxygen consumption associated with proton leak. We reported a trend towards an increase in basal respiratory rate as well as ATP-dependent respiration, and proton leak in AD-D as compared to AD-MCI and CTRL fibroblasts (Figure 3E, F and I, J). We noticed a tendency towards an increase in maximal respiratory rate in both AD-MCI and AD-D versus CTRL fibroblasts (Figure 3G) and an increase in the spare respiratory capacity, corresponding to the difference between the basal and the maximal respiration in AD-MCI as compared to CTRL and AD-D fibroblasts (Figure 3H). ECAR was also increased in AD-D versus CTRL fibroblasts at both baseline and under stress conditions (i.e. addition of oligomycin and FCCP) (Figure 3K, L). These data unexpectedly indicate a high metabolic activity in AD-D fibroblasts. Accordingly, we showed that glycolysis- and OXPHOS-mediated ATP production (GlycoATP and MitoATP respectively) were enhanced in AD-D fibroblasts (Figure 3M). Interestingly, correlation analyses showed that proton leak significantly correlates with CDR-SOB, MMSE and PiB-GCI (Suppl. Fig. 1G-I).

Altogether, these results support mitochondrial dysfunctions in AD-D fibroblasts and noticeable changes in AD-MCI fibroblasts as compared to CTRL. They also reveal a high metabolic activity in AD-D fibroblasts likely impacting mitochondrial fitness as manifested by enhanced mitochondrial potential and proton leak activity, this later being associated with AD cognitive hallmarks.

### 3.4- Mitophagy process is defective in AD-MCI and AD-D fibroblasts

After uncovering several mitochondrial dysfunctions in AD-MCI and AD-D fibroblasts, we studied mitophagy, the specific process of degradation of dysfunctional mitochondria. We first showed an increased colocalization of LC3-GFP probe with Mit-RFP probe, demonstrating that the number of mitophagosomes is slightly increased in AD-MCI versus CTRL reaching significance in AD-D fibroblasts (Figure 4A, B). We then found a significant increase in the localization of LAMP1-GFP probe with Mit-RFP probe in both AD-MCI and AD-D as compared to CTRL fibroblasts reflecting an enhanced number of mitolysosomes (Figure 4C, D). Together, these results would suggest enhanced mitophagy process in AD-MCI and AD-D fibroblasts. If this was the case, we would expect an enhanced degradation of mitochondria and a decrease in the level of mitochondrial proteins in AD fibroblasts versus CTRL. However, SDS-PAGE analyses using mitochondria-enriched fraction showed a significant accumulation of several mitochondrial proteins (TOMM20, CPN10 and CoxII) in AD-D fibroblasts (Figure 4E-H), correlating with MMSE (i.e. CoxII) and CDR-SOB (i.e. TOMM20) scores but not with PiB-GCI (Figure 4I). Moreover, by using Cox8-EGFP-mCherry mitophagy probe^23^, we observed a decrease in the localization of isolated mitochondria within lysosomes (number of red dots) in AD-MCI and AD-D fibroblasts as compared to CTRL (Fig 4J-L). This likely reflects a defective fusion of mitophagosomes with lysosomes and/or defective lysosomal function. Altogether, these data suggest a blockade of the mitophagy process and a defective degradation of dysfunctional mitochondria in AD-MCI and AD-D fibroblasts, that is pronounced in AD-D cells.

**Fig. 4.**
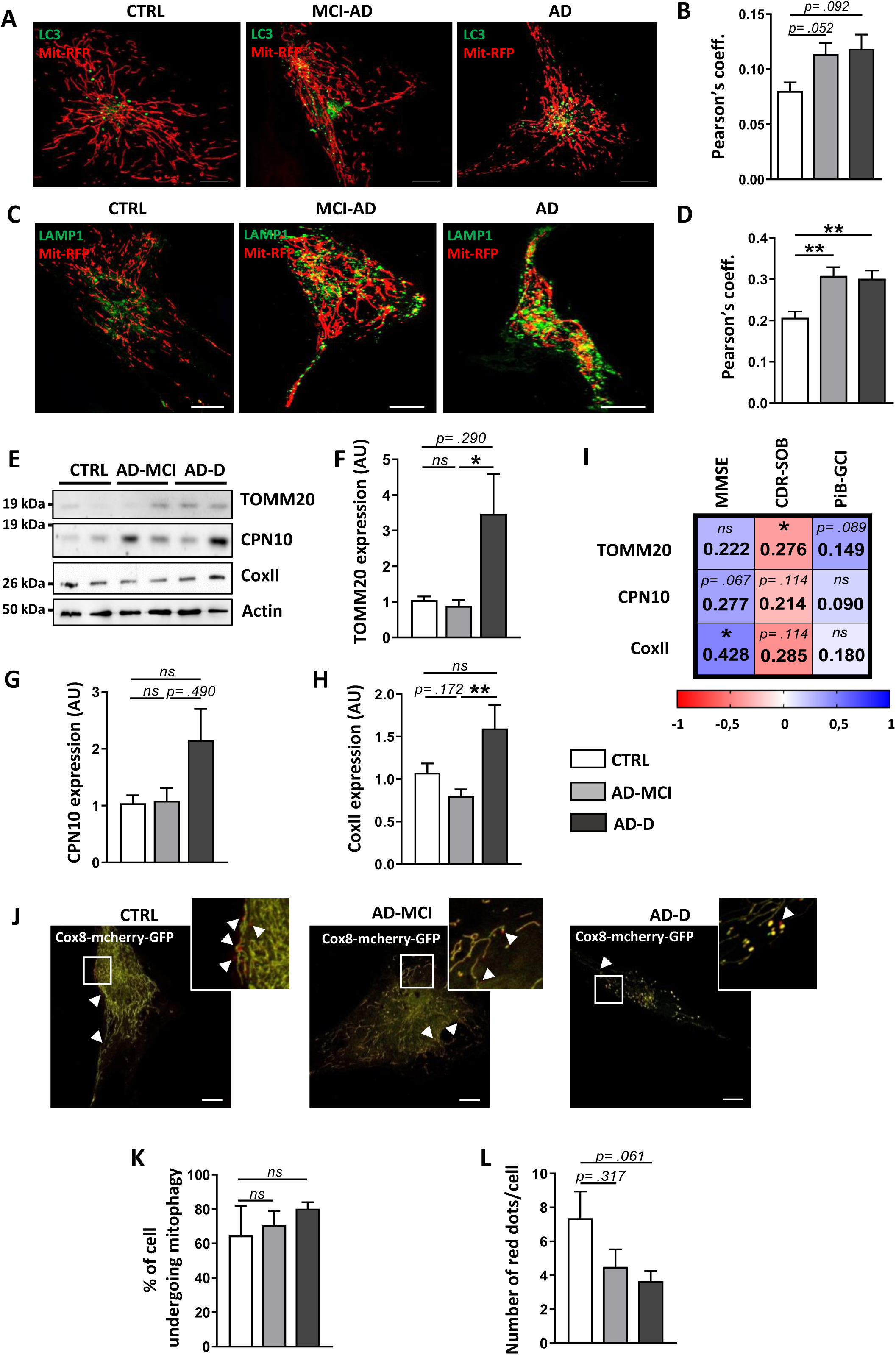
Mitophagosome, mitolysosomes and mitochondria content analyses reveal defective mitophagy process in AD-D fibroblasts. **A, C)** Representative images of Mit-RFP (red) LC3-GFP or LAMP1-GFP (green) signals in CTRL, AD-MCI and AD-D fibroblasts. Merge signal (yellow) reflects the colocalization of the red and the green signals. Scale bars = 2 μm. **B, D)** Quantitative graphs of the colocalization (Pearson’s coefficient) of Mit-RFP with LC3-GFP (n=5), AD-MCI (n=6), AD-D (n=5). (B) or of Mit-RFP with LAMP1-GFP (D). Means ± SEM obtained from CTRL (n=5), AD-MCI (n=5), AD-D (n=5). *p< .05; **p< .001; ns: not significant. **E-F)** Representative WB (E) and quantitative graphs of TOMM20 (F), CPN10 (G), and CoxII (F) expression levels ± SEM obtained in mitochondria-enriched fraction of CTRL (n=9), AD-MCI (n=11) and AD-D (n=9) fibroblasts. Data were normalized to CTRL fibroblasts. *p< .05; ns: not significant. **I)** Heat map of the correlation matrix computing linear regression (R²) value between TOMM20, CPN10 and CoxII expression levels and the clinical Dementia Rating (Sum Of Boxes) (CDR-SOB), the Mini Mental State Examination (MMSE) and the β-amyloid plaques burden assessed using the ^11^C-labelled Pittsburgh compound B (Global Cortical Index) (PiB-GCI) through positrons emission tomography (PET-scan) at patient’s inclusion. Color scale of 1 corresponds to the maximum positive correlation value (blue) and −1 corresponds to the maximum negative correlation value (red). Correlation analyses took into consideration the age of patients and the delay between the inclusion date and the date of skin biopsy as covariates. *p< .05; ns: not significant for every pair of data sets. **J)** Representative images of Cox8-EGFP-mCherry expression in CTRL, MCI-AD and AD-D fibroblasts. **K, L)** Quantitative graph of the percentage of cells undergoing mitophagy ± SEM (K), and of the number of red dots per cell ± SEM (L) obtained from CTRL (n=4), AD-MCI (n=5) and AD-D (n=4). ns: not significant.

### 3.5- Autophagy process is altered in AD-MCI and AD-D fibroblasts

In line with a mitophagy failure in AD fibroblasts, we hypothesized that the general autophagy is also altered in AD-derived fibroblasts. By using confocal microscopy, we quantified the number and size of LC3 and LAMP1 dots by respectively expressing LC3-GFP and LAMP1-GFP probes. Interestingly, we showed that LC3 dots number is significantly increased in AD-MCI fibroblasts as compared to CTRL and AD-D groups. In addition, we reported that LC3 size is significantly increased in AD-D fibroblasts as compared to CTRL and AD-MCI groups (Figure 5A-C) suggesting an enlargement of autophagosomes. Accordingly, we showed a significant decrease in the number of LAMP1 puncta in AD-D fibroblasts (Figure 5D, E) as well as an increase size of LAMP1 puncta in AD-MCI and AD-D fibroblasts as compared to CTRL (Figure 5D, F), thus highlighting swollen and potentially dysfunctional lysosomes.

**Fig. 5.**
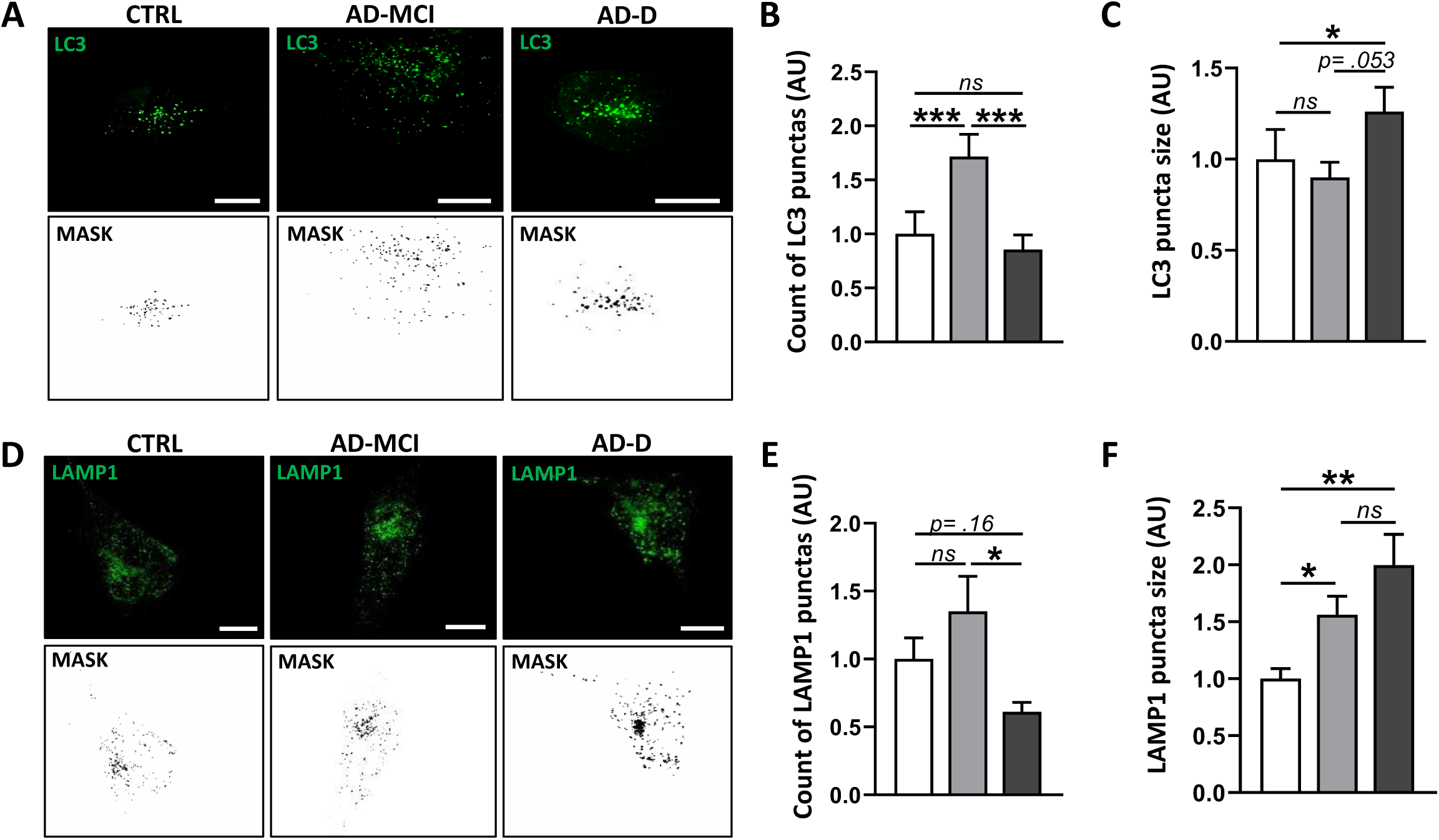
Autophagosomes and lysosomes analyses highlight a failure of autophagy in AD-D fibroblasts. **A, D)** Representative images and masks of LC3-GFP (A) and LAMP1-GFP (D) signals in CTRL, AD-MCI and AD-D fibroblasts. Scale bar =2 µm. **B, E)** Quantitative graph of the number ± SEM of LC3-GFP punctae (B) and of LAMP1 punctae (E). **C, F)** Quantitative graph of the size ± SEM of LC3-GFP punctae (C) and of LAMP1 punctae (F). Data were obtained in CTRL (n=5), AD-MCI (n=6) and AD-D (n=5) fibroblasts. *p< .05; **p<0.01; ***p<0,001; ns: not significant.

### 3.6- Lysosomal degradation is defective in AD-D fibroblasts and correlate with cognitive deficits

To assess whether the above-described LAMP1 alterations is linked to lysosomal defects, we measured lysosomal activity by using complementary approaches. We first used LysoTracker Red Probe to stain functional acidic lysosomes (pH 4) and revealed a significant decrease in LysoTracker intensity in AD-D versus CTRL and AD-MCI fibroblasts (Figure 6A, B). This demonstrates a defect of lysosomal acidification in AD-D fibroblasts. Accordingly, we confirmed that the degradation process of lysosomes is defective in AD-D fibroblasts as shown by a decrease of the cathepsin D (CTSD) lysosomal enzyme activity (Figure 6C-E). Interestingly, we revealed a significant negative correlation between CTSD activity and the CDR-SOB as well as a positive correlation with the MMSE (Figure 6F, G) while CTSD activity does not correlate with the PiB-GCI (Figure 6H). Together, these data (Figures 4 to 6) fully demonstrate a defective autophagy and mitophagy processes in AD fibroblasts linked to a failure of lysosomal degradation and correlating with cognitive deficits.

**Fig. 6.**
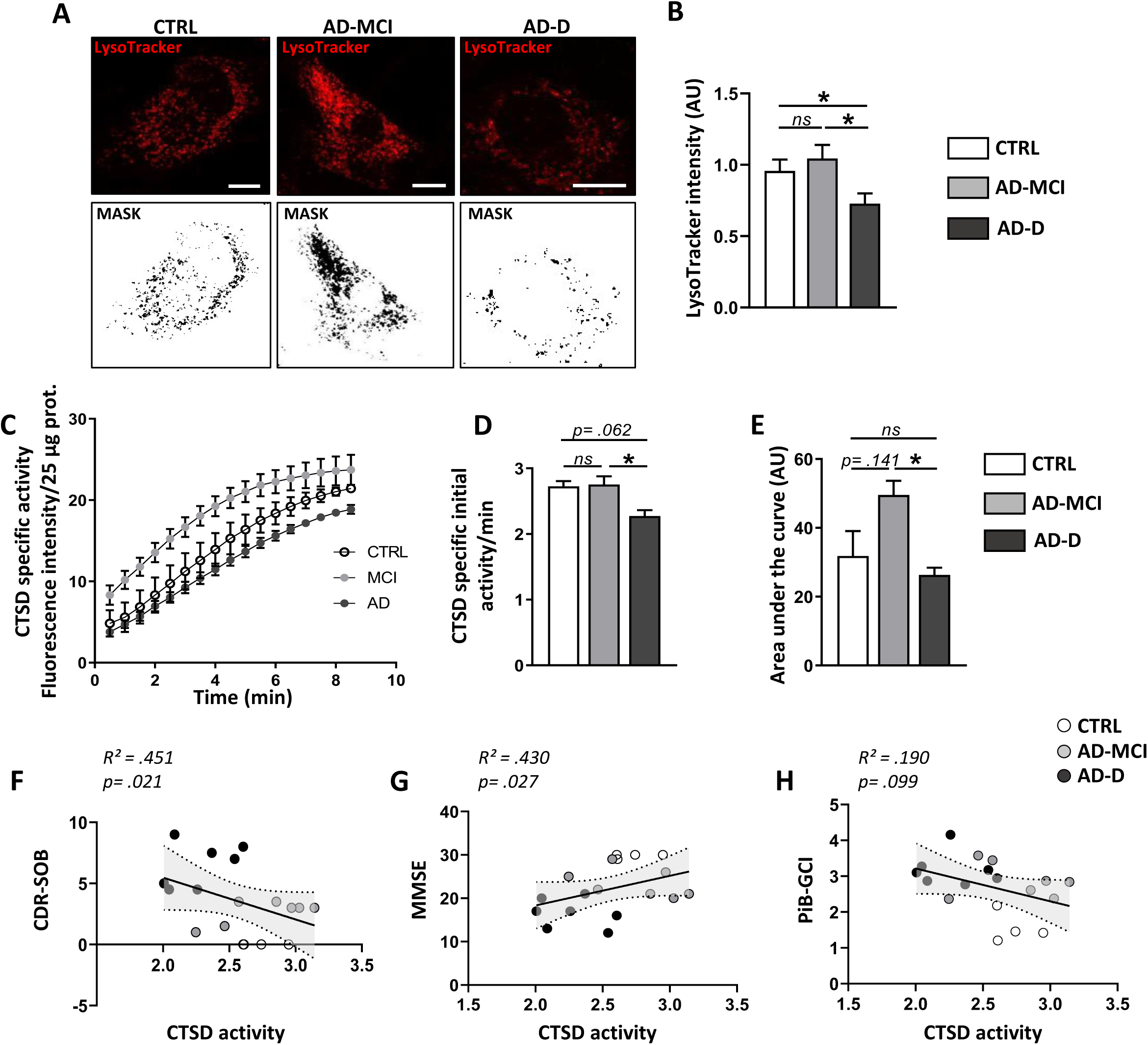
Lysosomal pH and cathepsin D activity analyses demonstrate defects of lysosomal degradation in AD-D fibroblasts. **A)** Representative images and masks of LysoTracker signal in CTRL, AD-MCI and AD-D fibroblasts. **B)** Quantitative graph of LysoTracker mean intensity ± SEM obtained from CTRL (n=4), AD-MCI (n=6) and AD-D (n=3) fibroblasts. Scale bar =2 µm. **C-E)** Cathepsin D (CTSD) mean specific activity ± SEM (C) and quantification of the initial specific activity ± SEM (D) and the area under the curve ± SEM (E) obtained from CTRL (n=4), AD-MCI (n=7) and AD-D (n=7). *p< .05; **p< .01. **F-I)** Correlation plots between CTSD activity and the Clinical Dementia Rating (Sum Of Boxes) (CDR-SOB) (F), Mini-Mental State Examination (MMSE) of patients (G) and, the β-amyloid plaques burden assessed using the ^11^C-labelled Pittsburgh compound B (Global Cortical Index) (PiB-GCI) through positrons emission tomography (PET-scan) (H), at patient’s inclusion including CTRL (white dots), AD-MCI (grey dots) and AD-D (black dots). The linear regression was used to determine goodness of fit (R^2^) value and *p* values for the correlation of every pair of data sets taking into consideration the age of patients and the delay between the inclusion date and the date of skin biopsy as covariates.

### 3.7- APP and CTFs accumulation in mitochondria of AD-D fibroblasts correlates with cognitive deficits, mitochondrial proteins load and reduced cathepsin D activity

Several studies reported that full-length APP is also localized^28,29^ and processed in mitochondria^30–32^. Accordingly, APP metabolites (C99, AICD and Aβ) accumulate in mitochondria-associated membranes^33,34^. Furthermore, our team and others recently reported that mitochondria structure and function defaults as well as mitophagy failure are triggered by APP-CTFs accumulation within mitochondria^5,35^. Hence, we sought here to analyze the expression levels of full-length APP and APP-CTFs (C99, C83 and AICD) in mitochondrial fraction of fibroblasts from CTRL, AD-MCI and AD-D patients. SDS-PAGE analyses showed an accumulation of full-length APP and a trend towards an increase of the levels of APP-CTFs in AD-D as compared to CTRL and AD-MCI fibroblasts (Figure 7A, B). Interestingly, linear regression analyses revealed statistically significant positive correlations between the expression levels of full-length APP, C83 and C99 fragments with Aβ load measured by the PiB-GCI (Figure 7C). We also highlighted a negative correlation of C99 expression levels, but not APP and C83, with the MMSE score (Figure 7C). In addition, while the accumulation of APP-CTFs did not correlate neither with the mitochondrial structure (Suppl. Fig. 2A, B) nor with the mitochondrial function (Suppl. Fig. 2C-E), we interestingly unveiled a positive correlation between the accumulation of APP-CTFs with the accumulation of the mitochondrial proteins TOMM20 (i.e. C99 and C83, with a higher significance for C99) and CPN10 (i.e. C99) and showed a negative correlation between the accumulation of APP-CTFs and the CTSD activity (i.e. C99 and C83, with a significance for C99) (Figure 7D-F). Altogether, these data suggest an accumulation of APP and APP-CTFs in mitochondria of AD-D fibroblasts which is closely related to cognitive deficit and lysosomal defect occurring in AD-D patients.

**Fig. 7.**
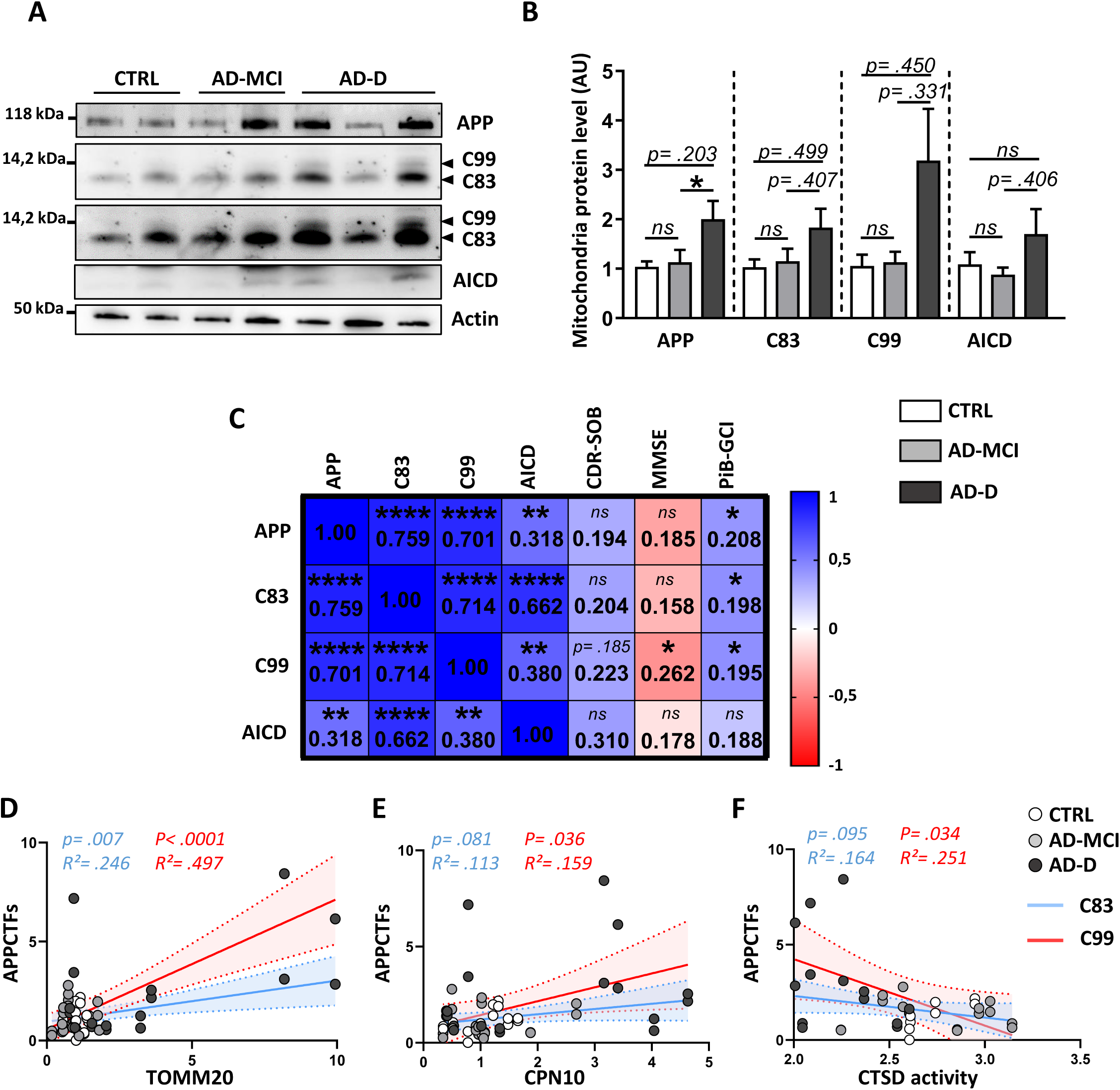
APP and APP-CTFs accumulation in the mitochondria of AD-D fibroblasts correlates with cognitive impairments, mitochondrial protein load and CTSD reduced activity. **A, B)** Representative WB (A) and quantitative graphs of the expression levels ± SEM (B) of APP full-length (APP Total) and APP-CTFs (C99, C83 and AICD) obtained in mitochondria-enriched fraction of CTRL (n=9), AD-MCI (n=11) and AD-D (n=9) fibroblasts. Data were normalized to CTRL fibroblasts. *p< .05. **C)** Heat map of the correlation matrix computing the linear regression (R²) between protein levels of APP full-length, C83, C99 and AICD fragments with the clinical Dementia Rating (Sum Of Boxes) (CDR-SOB), the Mini Mental State Examination (MMSE) and the β-amyloid plaques burden assessed using the ^11^C-labelled Pittsburgh compound B (Global Cortical Index) (PiB-GCI) through positrons emission tomography (PET-scan) at patient’s inclusion. Color scale of 1 corresponds to the maximum positive correlation value (blue) and −1 corresponds to the maximum negative correlation value (red). Correlation analyses took into consideration the age of patients and the delay between the date of inclusion and the date of skin biopsy as covariates. *p< .05; **p< .01; ****p< .0001; ns: no significant, for the correlation of every pair of data sets. **D-F)** Correlation plots between APP-CTFs and TOMM20 (D), CPN10 (E), or CTSD activity (F) including CTRL (white dots), AD-MCI (grey dots) and AD-D (black dots).

## 4 DISCUSSION

Several clinical studies suggest that fibroblasts obtained from AD patients harbor cellular dysfunctions observed in post-mortem AD brains^20^. In this study, we investigated mitochondria structure and function and examined autophagy and mitophagy processes in fibroblasts obtained from AD patients with a final objective to identify potential correlations between changes in mitochondria homeostasis and AD clinical onset.

First, we report an alteration of mitochondrial ultrastructure (Figure 1) and a decrease in mitochondrial volume and the average branch-length in AD-D fibroblasts as compared to AD-MCI and controls (Figure 2). These results are in line with previous studies conducted in distinct SAD-derived fibroblasts^12,13,36,37^. In addition, other studies have shown that fibroblasts from FAD patients also exhibit a reduction in mitochondrial number^15,38^. Together, these studies, apart from disclosing that alterations of the integrity and of the morphology of the mitochondrial network are commonly observed in SAD and FAD fibroblasts, they agree well with the data reporting fragmented mitochondria in post-mortem AD brain samples^39^. Mitochondria phenotypic changes are dependent on the balance between mitochondrial fusion and fission processes that are governed by Mitofusin 1 and 2 (MFN1, MFN2) forming homotypic and heterotypic interactions with OPA1 protein to initiate mitochondria fusion, and by FIS1 protein which interacts with dynamin-related protein1 (DRP1) at mitochondrial fission sites^40^. Interestingly, Wang *et al.* reported altered expression of mitochondria fusion and fission proteins in cortical samples from AD patient’s brains supporting enhanced mitochondria fission^39^. Likewise, Manczak *et al.*, showed that the mRNA and protein levels of FIS1 and DRP1 are increased in the frontal cortex of patients with early, definite, and severe AD^41^. However, mitochondrial structure phenotypes alterations in AD fibroblasts are not corroborated by consistent modulation of mitochondrial fission and fusion proteins expression. Pérez *et al.,* showed increased MFN1 expression level in SAD fibroblasts^12^ suggesting a compensatory mechanism as mitochondria hyper fusion may protects mitochondria from excessive mitophagy during stress conditions. A recent study by Drabik *et al.,* demonstrated a drop in the frequency of fusion-fission events in SAD fibroblasts, associated with a reduction in both fission and fusion proteins^25^.

Mitochondria morphology finely shapes organelle function and vice-versa. Hence, both mitochondrial membrane depolarization or hyperpolarization are considered as key features of mitochondria damage^42,43^. Furthermore, while controlled production of mitROS participates to mitochondria-nucleus communication ^44^, an excess of mitROS production is deleterious as it leads to oxidative damage of mitochondrial and cellular proteins, lipids and DNA^45^. We unravel here an increase in mitROS levels under basal conditions in AD-MCI but not in AD-D fibroblasts (Figure 3A). This observation does not corroborate the results of other groups describing an increase in mitROS in SAD^12,46–48^ and FAD fibroblasts^49^. However, it is important to emphasize that the stratification of AD-MCI and AD-D is significantly different between our cohort and when indicated in the above cited studies. Indeed, in our study, AD patients were stratified according to strict clinico-biological criteria, including the positivity of CSF biomarkers and amyloid PET imaging. Hence, MMSE mean score of AD-MCI patients was quantified as 24.3±2.97 and that of AD-D patients’ as 16.5±3.36 (Table 1). However, the MMSE mean score of AD patients in the study of Drabik *et al.,* ranged between 20 and 23, i.e. similar to that of our AD-MCI group^46^. Therefore, enhanced mitROS observed in AD-MCI fibroblasts in our cohort is likely consistent with the results previously reported in different AD fibroblasts cohorts. Moreover, even if AD-D fibroblasts mitROS level is comparable to that observed in CTRL fibroblast under basal condition, it is dramatically and significantly increased under stress condition (Figure 3A, B). Together, these results demonstrate that mitROS is commonly enhanced in SAD and FAD fibroblasts, likely that this occurr at early disease stages. Our data unveil that basal- and ATP-linked respiration is enhanced in AD-D fibroblasts and that the spare respiratory capacity, reflecting mitochondrial reserve under stress condition is increased in AD-MCI fibroblasts. These data point to specific metabolism signatures in AD-MCI and AD-D fibroblasts revealing on the one hand a higher metabolism capacity in AD-D fibroblasts and on the other hand a higher adaptive capacity of AD-MCI fibroblasts to stress condition. In addition, we also show that proton leak is enhanced in AD-D fibroblasts and is corroborated by the significant mitochondrial membrane hyperpolarization as compared to AD-MCI and CTRL fibroblasts (Figure 3C, D). Mechanistically, proton leak is operated by uncoupling proteins and is linked to the inhibition of the ATPase F_1_F_0_ activity or its operation in a reverse mode^42^. It is well known, that the activation of proton leak pathways under physiological conditions minimize oxidative damage by tempering mitochondrial potential and mitochondrial superoxide production^50,51^. However, when proton leak is installed chronically, it may impair the adaptability of cells to stress conditions, as observed in AD fibroblasts.

Mitochondrial fragmentation and increase in mitROS production are major features linked to mitophagy^52^. Our data indicate enhanced mitophagosomes and mitolysosomes formation in AD-D versus CTRL fibroblasts (Figure 4A-D), supporting mitophagy initiation in AD. However, we observed an enlargement of autophagosome and lysosomes indicative for defective degradation processes (Figure 5A-F). Previous studies have reported an enlargement of endosomes in both PBMCs and fibroblasts of IMABio3 cohort^17,20^. As a whole, these results demonstrate a global alteration of the endolysosomal compartments in SAD and FAD fibroblasts as it was already described^53–55^. AD-D fibroblasts display a significant increase in mitochondrial resident proteins levels (i.e. TOMM20, CoxII as well as CPN10) as compared to CTRL cells (Figure 4E). This is expectedly linked to mitophagy defect. Accordingly, an increase in TOMM20 protein level^53^ and a mitophagy failure molecular signature were also described in SAD brains^5^.

The enlargement of lysosomes in AD-D fibroblasts support a defect of their activity that was linked to reduced acidification and decreased CTSD protease activity (Figure 6 A-E). We also reveal a correlation of CTSD activity with cognitive decline (Figure 6 F, G). Interestingly, genome wide-association study uncovered *CTSD* and cathepsin B (*CTSB*) genes as potential genetic risk factors for AD^56,57^. Together, these data suggest that the failure of mitophagy/autophagy in AD-D fibroblasts is linked to lysosomal defects and may constitute a reliable indicator of AD progression in peripheral cells.

APP-CTFs accumulate in the brains of AD patients^5,58^ and may serve as a potential biomarker for AD^59^. We report for the first-time, increased APP and APP-CTFs (C83, C99 and AICD) protein levels in the mitochondrial fractions in AD fibroblasts correlating with AD-related neuropsychological scores (Figure 7). Our study and others demonstrate that APP-CTFs accumulation trigger mitochondria structure and function default as well as mitophagy failure^5,32,35^. In parallel, altered mitochondrial fitness and homeostasis could also affect APP processing as well as Aβ production leading to a vicious circle in the pathophysiological process of AD^60^. Our study showed a positive correlation of APP-CTFs accumulation with mitochondrial proteins levels but not with mitochondrial structure and function alterations. These data led us to assume that peripheral mitochondrial alterations observed in AD patients are not exclusively linked to APP-CTFs accumulation. Indeed, many mitochondrial alterations accompanying the pathophysiological development of SAD were suggested to be driven by genetic risk factors^61^. More particularly, some mitochondrial genes were shown to be associated with SAD^56^, including *TOMM40* located within the locus of *APOE* genetic risk factor^62^. TOMM40 is a component of the cation-selective translocation pore of the outer membrane translocase (TOM) complex allowing the entry of mitochondrial proteins encoded by nuclear genes^63^. It is thus plausible that *TOMM40* genetic variants and/or alterations of the expression of other members of TOM complex such as TOMM20 (i.e as observed in AD-D samples) compromise mitochondrial protein translocation and subsequently function^62^. Furthermore, APP, Aβ peptides and likely APP-CTFs interact and/or are imported into mitochondria through TOM complex^29,64^. Polymorphisms in *TFAM* gene, encoding the mitochondrial transcription factor A (TFAM) protein was also associated with late-onset AD risk^56,65,66^. TFAM plays a key role in the maintenance of mtDNA integrity by regulating mitochondrial transcription and mtDNA copy number. Importantly, TFAM expression is reduced in both MCI and AD PBMCs and correlates with AD severity and mtDNA content^67^. Polymorphisms in *COX7C* gene, encoding for COX7C protein a component of OXPHOS complex IV, was recently described in the IMABio3 cohort^20^. Moreover, COX7C has lower expression in the blood of MCI/AD relative to age-matched controls^68^. Lastly, a new variant of a mitochondrial microprotein called SHMOOSE (*SHMOOSE.D47N*), is associated with AD risk and rapid cognitive decline, and correlates with reduced cortical volume^69^. SHMOOSE increases oxygen consumption rate, boost mitochondrial spare respiratory capacity and bind the mitofilin protein in the mitochondrial inner membrane, known to regulate cristae junctions^69^. Interestingly, SHMOOSE-related functions are altered in IMABio3 AD fibroblasts.

To our knowledge, this is the first study reporting mitochondrial ultrastructure, morphology, function alterations and mitophagy dysfunctions in fibroblasts obtained from a clinically characterized SAD cohort including AD-MCI and AD-D patients. This study unravels that the observed mitochondrial alterations mostly recapitulate those reported in AD brain samples. Accurate AD diagnosis is still needed; thus, it is essential to uncover new peripheral biomarkers for the identification of early changes in peripheral cells/fluids in AD patients. Our study provides evidences that mitochondrial alterations and defect in lysosomal degradation in peripheral tissues are potential biomarkers accompanying the progression of AD. We also report for the first time an accumulation of APP-CTFs in mitochondria in AD fibroblasts. Therefore, peripheral APP-CTFs levels might also be considered as an additional pathophysiological marker of AD.

All over, this study opens perspectives to investigate the molecular mechanisms underlying mitochondrial dysfunction in peripheral AD cells.

## ACKNOWLEDGMENTS

We acknowledge members of the clinical IMABio3 team: Dr Amer Alnajjar-Carpentier, Consultation mémoire, Hôpital d’Orsay, Orsay, France; Dr Michel Logak, Service de Neurologie, Hôpital Saint Joseph, Paris, France; Dr Sara Leder and Dr Dominique Marchal, Consultation mémoire, Institut des Invalides, Paris, France; Dr Hélène Pitti-Ferandi, Consultation mémoire, Clinique de la Porte Verte, Versailles, France; Dr Hélene Brugeilles, Service de Neurologie, Hôpital Mignot, Versailles, France; Dr Brigitte Roualdes, Service de Neurologie, Hopital Henri Mondor, Versailles, France; Dr Agnes Michon, Consultation mémoire, Hôpital de la Salpêtrière, Paris, France. We wish to thank the DNA and Cell Bank platform from ICM for producing the first vials of fibroblasts.

## CONSENT STATEMENT

IMABio3 and Shatau7-Imatau study cohorts were approved by a French Ethics Committee (NCT01775696 and NCT02576821-EudraCT2015-000257-20). All subjects provided written informed consent prior to participating.

## AUTHOR CONTRIBUTIONS

MC conceived and designed the study. FE designed the experiments, conducted and analysed the experiments. MC and FE analysed data and wrote the manuscript. PFK participated to lysosomal and CTSD activities. SLG conducted electron microscopy experiments. LX, MCP and MS provided primary cultures of fibroblasts. JL, MS, M CP discussed data. JL, MB, GD and MS collected patient data. MC, FE, JL, MS, M CP and FC read and edited the manuscript.

