## Supplementary material for "Mitochondria alterations in fibroblasts from sporadic Alzheimer’s disease (AD) patients correlate with AD-related clinical hallmarks": Supplementary materials.docx

**Suppl. Fig. 1. Correlation analyses between the mitochondrial function and the patients’ clinical data. A-F)** Correlation plots between TMRM (A-C) and MitoSox (D-F) and the CDR-SOB (A, D), MMSE (B, E), and PiB-GCI (C, F) at patient’s inclusion including CTRL (white dots), AD-MCI (grey dots) and AD-D (black dots). The linear regressions were used to determine *P* and goodness of fit (R^2^) values. **D-F)** Correlation plots between the proton leak and the CDR-SOB (G), MMSE (H), and PiB-GCI (I) at patient’s inclusion including CTRL (white dots), AD-MCI (grey dots) and AD-D (black dots). The linear regression was used to determine goodness of fit (R^2^) values.

**Suppl. Fig. 2. Correlation analyses between the accumulation of APP-CTFs and mitochondria structure or function**. **A-E)** Correlation plots between APP-CTFs and mitochondria classes I (A) and IV (B), TMRM mean intensity (C), MitoSox mean intensity (D), and proton leak (E). The linear regression with C83 (blue) or C99 (red) were used to determine *P* and goodness of fit (R^2^) values.
