## Supplementary figures and images for "Mitochondria alterations in fibroblasts from sporadic Alzheimer’s disease (AD) patients correlate with AD-related clinical hallmarks"

### Supplementary Figure 1.TIF

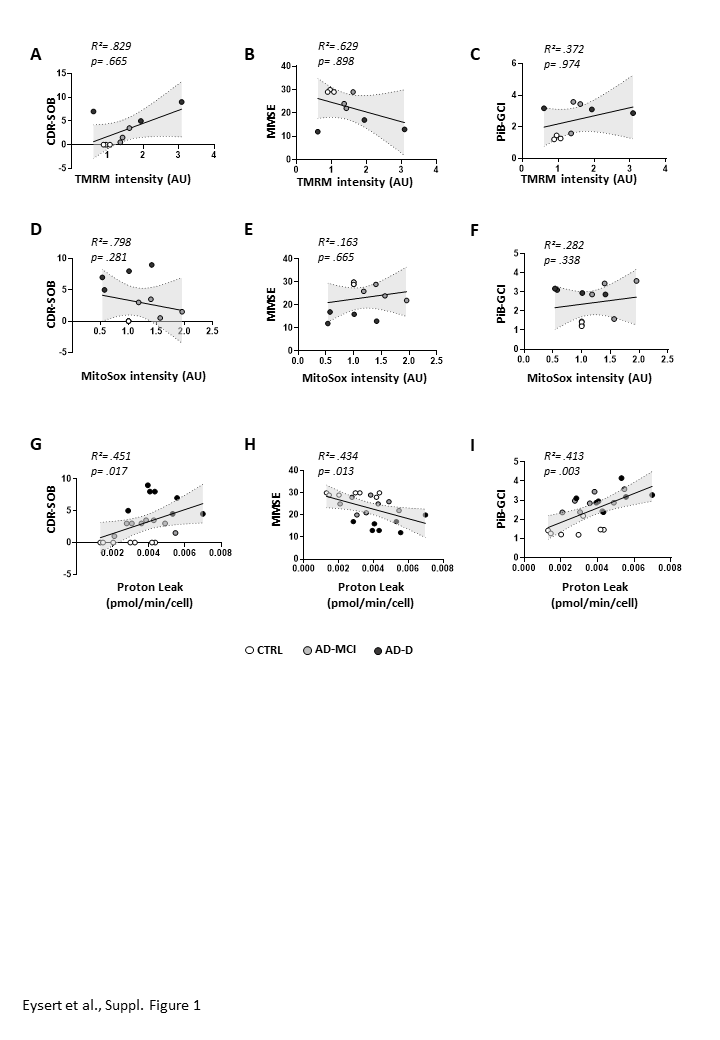

### Supplementary Figure 2.TIF

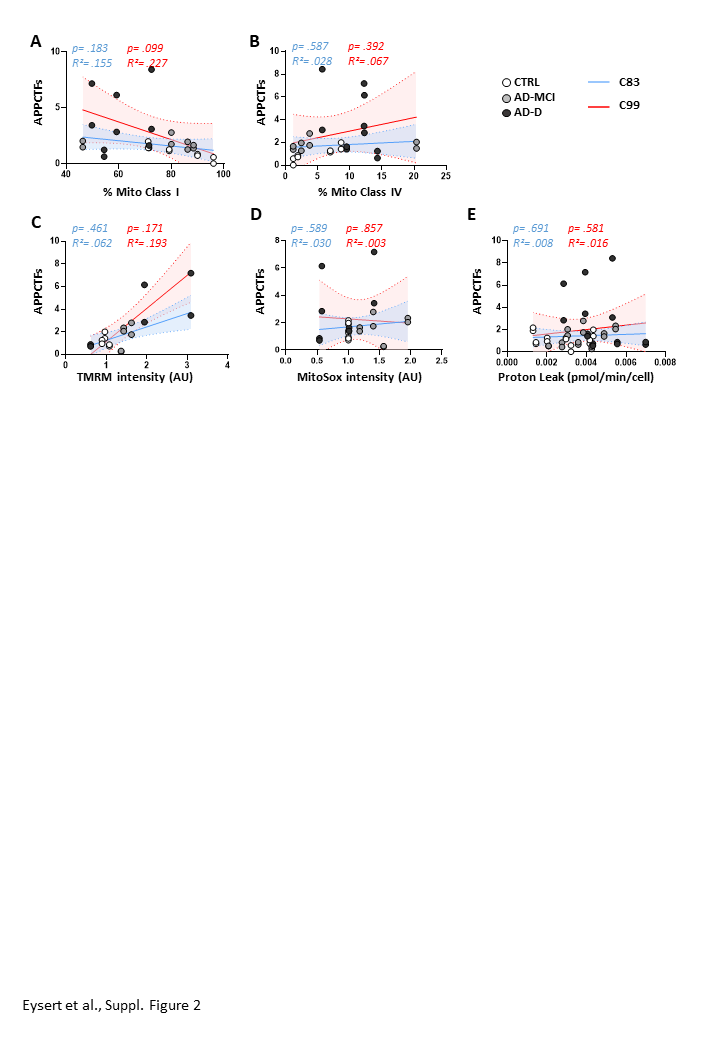
